# Cyclical Regression Covariates remove the major confounding effect of cyclical developmental gene expression with strain-specific drug response in the malaria parasite *Plasmodium falciparum*

**DOI:** 10.1101/2020.04.17.044552

**Authors:** Gabriel J. Foster, Katrina Button-Simons, Katelyn Vendrely, Jeanne Romero-Severson, Michael T. Ferdig

**Author notes:** Author Information: Gabriel J. Foster, Katrina Button-Simons, Katelyn Vendrely, Jeanne Romero-Severson, Michael T. Ferdig.

## Abstract

**Background:** The cyclical nature of parasite gene expression in the intraerythrocytic development cycle (IDC) in human blood confounds the accurate detection of specific transcriptional differences due to drug resistance in *Plasmodium falciparum*. Here, we propose the use of cyclical regression covariates to eliminate the major confounding of developmentally driven transcriptional changes with changes due to drug response. We show that elimination of this confounding can reduce both Type I and Type II errors, and demonstrate the effect of approach on real data.

**Results:** We apply this method to two publicly available datasets, and demonstrate its ability to reduce the potential confounding of differences in expression due the species-specific intraerythrocytic development cycle from strain-specific differences in drug response. We show that the application of cyclical regression covariates has minimal impact on the pool of transcripts identified as significantly different in a dataset generated from single timepoint clinical blood samples with low variance for developmental stage and a profound impact on another clinical data set with more variance among the samples for developmental stage.

**Conclusions:** Cyclical regression covariates have immediate application to studies where *in-vitro* synchronization of all samples to the same developmental timepoint is not feasible, primarily parasite transcriptome sequencing direct from clinical blood samples, a widely used approach to frontline detection of emerging drug resistance.

## Background

The *P. falciparum* parasite causes the most lethal form of malaria, a vector-borne disease that killed an estimated 405,000 people in 2018, 272,000 of them children under the age of five [1]. Although malaria prevention strategies and early treatment efforts have reduced the worldwide incidence of malaria 18% since 2010, the persistence of malaria is due in part to the rapid emergence and spread of parasites resistant to antimalarial drugs, including artemisinin-based combination therapies, the last line of defense in regions where multiple drug resistance has arisen [2-4].

Most antimalarial drugs, including artemisinin-based combination therapies (ACT) therapies, target the parasite in the intraerythrocytic development cycle (IDC), the blood stage initiated when the parasite invades red blood cells [5]. Whole transcriptome gene expression of the malaria parasite in the IDC, in combination with whole genome DNA sequence, can detect mutations and identify pathways that result in drug resistance [6-11]. However, comparative analysis of gene expression in drug susceptible and drug resistant strains remains a challenge due to the confounding effects of the cyclical gene expression that drives the *P. falciparum* development through the three asexual stages of IDC [12]. During the IDC, the vast majority of *P. falciparum* transcripts are expressed in a single sinusoidal pattern smoothly extending in a continuous cascade across a morphological progression through three asexual forms: ring, trophozoite, and schizont [13-15]. The period of the curve corresponds to one complete progression through the IDC [16]. This pattern is seen in other *Plasmodium* species and another apicomplexan parasite, *Toxoplasma gondii* [17]. Accurate detection of transcriptional differences between parasite strains due to drug resistance in the IDC requires that the comparison is made when both strains are at the same point in the IDC.

Single timepoint *in-vitro* transcriptional studies in *P. falciparum* typically rely on microscopy to identify the point at which half of an experimentally well-synchronized culture has converted from mature schizonts to new rings, the first stage of progression through the IDC, i.e. 0 hours post-invasion (hpi). [18-20]; samples are then taken a set number of hours after 0 hpi, based on the experimental parameters. In the case of *ex-vivo* studies, the parasites taken straight from patients are not laboratory cultured and thus *ex-vivo* samples are not experimentally synchronized with each other. Simple morphological examination of samples based on microscopy is not accurate enough to ensure that ex-vivo transcriptomes are appropriately aligned prior to differential expression analysis. [21].

In a recent large *P. falciparum* study, Mok et al. used k-means clustering on 1043 *ex-vivo* whole transcriptome samples to identify subgroups for study [22]; they defined three major subgroups. Post-hoc analysis staged the whole transcriptomes against the multi-timepoint transcriptome data of 3D7, where the authors found that the subgroups largely segregated by IDC progression. As progression through the asexual forms is a continuous rather a discrete process, IDC progression differences within these clusters could still lead to mis-attributing stage-based transcriptional differences to the variable under study (e.g. drug resistance). Another study applied a novel correction method to the Mok et al. data by removing the effect of a linear and polynomial covariate of IDC timepoint before differential expression analysis [23]. As segments of the broadly sinusoidal progression curve can be approximated by linear and polynomial functions, this method provides improved accuracy in sample sets with small variance in IDC progression. However, as the method does not specifically incorporate the cyclic nature of expression, it is prone to potential overfitting and is not useful for samples whose developmental stage distribution spans reinvasion of new uninfected RBCs.

Cyclical regression covariates are applied to wide range of biological data with cyclical covariance, including diarrheal severity, transmission of malaria, and seasonal changes in human gene expression [24-29]. Here we examine the use of cyclical regression covariates to disentangle expression differences due to developmental stage progression from differences in response to artemisinin in *P. falciparum* parasites. We demonstrate how small differences in developmental stage between two strains can generate both false positive and false negative associations between transcript level and response to treatment with the drug artemisinin, and show how these errors can be significantly reduced by using cyclical regression coefficients to align developmental progression in two different strains. We then apply this method to the Mok et al. set of *ex-vivo P. falciparum* transcriptional samples previously assessed by transcriptional staging against 3D7 without cyclical regression covariates, then grouped into three using k-means clustering [22].

## Results

### Cyclical regression covariates reduce both false positive and false negative associations

In the simple model, when a transcriptional sample is taken at the same point in developmental progression from two different *P. falciparum* strains with identical gene expression profiles, a lack of gene expression differential is properly observed (Figure 1A). However, when these same strains are (inadvertently) sampled at different points in their respective cyclical transcriptional progressions and compared without adjustment, a significant gene expression difference is falsely detected, a Type I error (Figure 1B). Two strains with a true gene expression difference sampled at the same point in IDC progression will show a significant difference that actually exists (Figure 1C). Sampling these same two strains at different timepoints in their respective progressions can generate the appearance of a difference where none exists (a Type I error) and obscure the true difference, a Type II error (Figure 1D).

**Figure 1:**
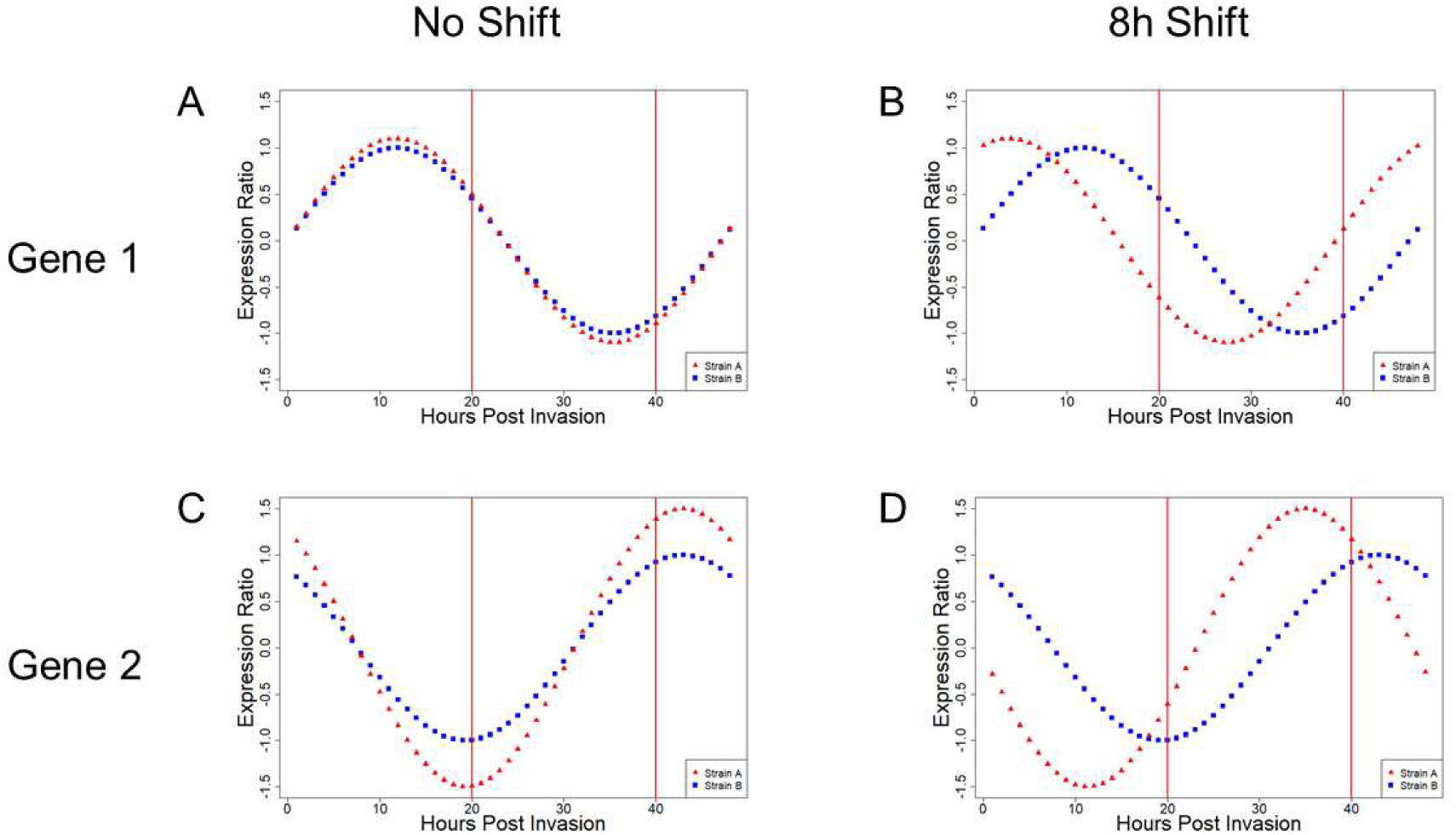
Errors caused by comparison of improperly aligned cyclical transcription data. Figure 1A and 1B show two model features with identical expression patterns. Figure 1A: When 0hpi is correctly aligned for both, single observations (red bars) correctly detect no difference. Figure 1B: When 0hpi is improperly aligned, single observations (red bars) detect a difference between strains where none exists. Figures 1C and 1D represent two model features with real differences in gene expression. Figure 1C: Proper alignment of 0hpi allows single observations (red bars) to detect the true difference. Figure 1D: Improper alignment of 0hpi obscures the true difference in single observations.

Examination of the transcriptional profiles of two features (PF3D7_1034400 and PF3D7_0926700) in two different *P. falciparum* strains (HB3 and 3D7) illustrates the effect of misalignment in real data [16, 30]. The sequence PF3D7_1034400 displays a characteristic wave pattern of expression across the entire IDC that is similar for both strains when transcriptional profiles correctly begin at 0hpi (Figure 2A). A computational shift of the sequence from HB3 eight hours forward results in the appearance of different expression between the two sequences across multiple timepoints, false positives driven solely by misalignment (Figure 2B). In the alternative case, sequence PF3D7_0926700 shows clear transcriptional differences at 40hpi between HB3 and 3D7 when t = 0hpi is correctly determined for both parasite strains (Figure 2C). Computationally shifting HB3 eight hours forward renders this transcriptional difference at 40hpi undetectable (a Type I error) and generates a false difference at 20hpi (a Type II error) (Figure 2D).

**Figure 2:**
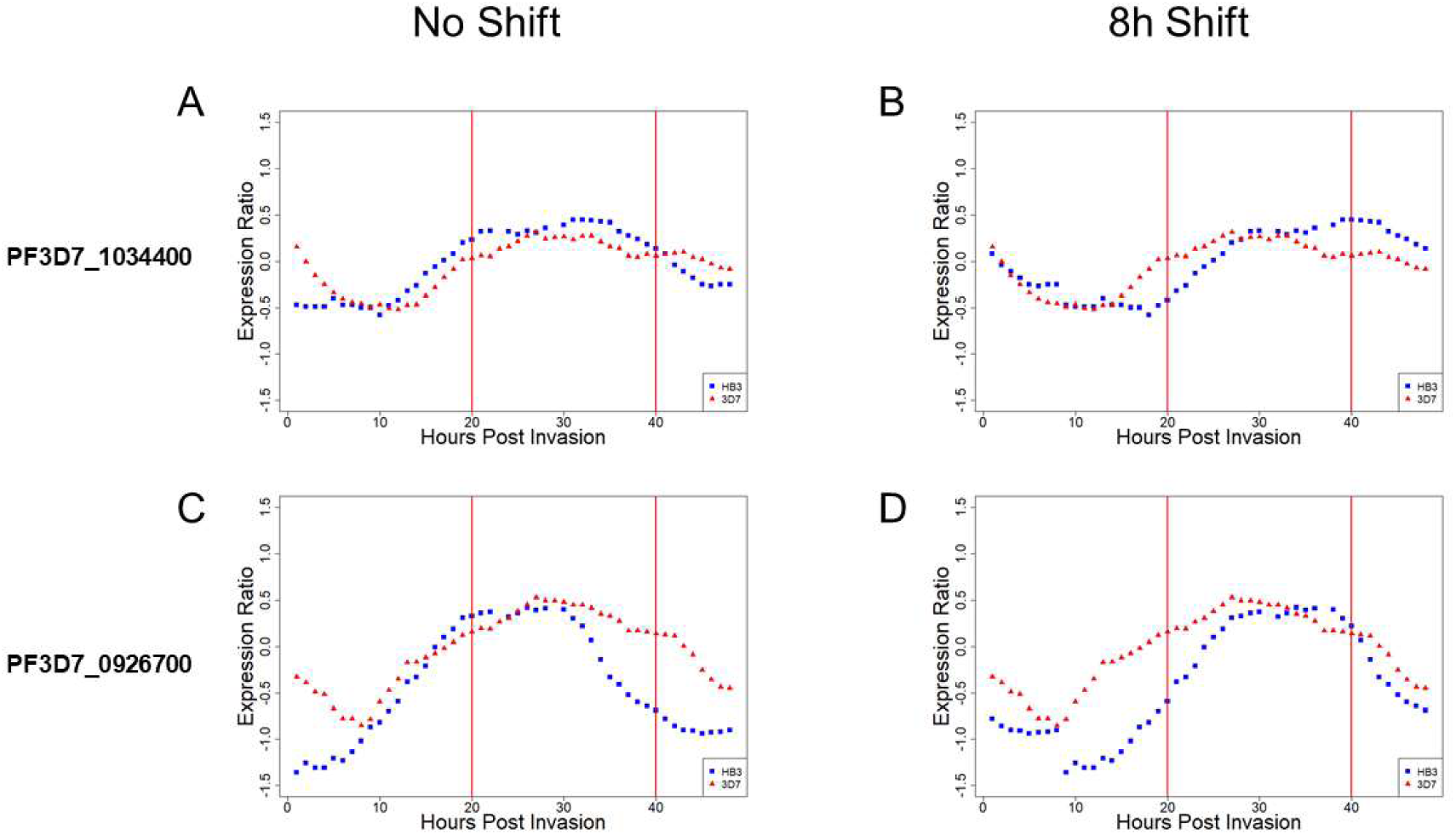
Comparison of *P. falciparum* transcriptional profiles of two features in strains HB3 and 3D7 data. 2A: Correct assignment of 0hpi shows little difference in the expression of PF3D7_1034400 between HB3 and 3D7. 2B: An 8 h shift in HB3 IDC progression timing produces differential expression at 20 h and 40 h due to the progression shift. 2C: In PF3D7_0926700, a genuine expression difference exists between HB3 and 3D7 at 40hpi. 2D: An 8 h shift in HB3 expression obscures the differential expression at 40 h, and generates a difference at 20 h; the misalignment creates both Type I and Type II errors. Figures 2A, 2B, 2C, and 2D are derived from transcriptional time course data from HB3 and 3D7[16, 30].

### Cyclical regression covariates remove the effect of stage progression on transcription in an entire dataset

We then tested the effect of cyclical regression covariates on a published *P. falciparum* dataset containing transcriptional profiles of the parasite strain (PB58) collected at three timepoints (6hpi, 26hpi, and 38hpi) with three biological replicates for each timepoint [31]. As the samples all were taken from the same parasite and the data has replication, any differential expression detected must solely be a result of different IDC progression times. Pairwise comparison of different timepoints (e.g. 6hpi with 26hpi) reveals the degree of false positive associations possible when the confounding effect of IDC progression is not removed (Table 1A). The large number of genes considered significantly expressed only as a result of progression differences demonstrates that any stage-blind analysis will be largely confounded by the effects of stage progression on transcription.

**Table 1:** Removing the effect of stage removes Type I errors. Three replications of whole transcriptome samples from PB58 at 3 timepoints; 6hpi, 26hpi, and 38hpi. A: In silico misalignment of the timepoints reveals the extent of expression differences that appear due to this misalignment (p < 0.01, FDR < 0.05). B: Application of cyclical regression covariates to correctly align the replications results in no differences in gene expression, the expected result of replication for a single strain in the haploid IDC stage.

|  |  | PB58 |  |  |
| --- | --- | --- | --- | --- |
|  |  | 6 | 26 | 38 |
| PB58 | 6 | - | 2989 | 294 |
|  | 26 | 2989 | - | 457 |
|  | 38 | 294 | 457 | - |

|  |  | PB58 |  |  |
| --- | --- | --- | --- | --- |
|  |  | 6 | 26 | 38 |
| PB58 | 6 | - | 0 | 0 |
|  | 26 | 0 | - | 0 |
|  | 38 | 0 | 0 | - |

We then performed a differential expression analysis using a linear model including the covariates (Methods). Briefly, if we consider IDC progression as a circle, we can calculate a sample’s location on that circle using its IDC time and include those coordinates as covariates in a linear model. After accounting for stage progression using cyclical regression covariates, our analysis finds no differentially expressed genes. (Table 1B) The method eliminated the confounding that caused the false positives in this dataset.

### Application of cyclical regression covariates to a set of ex-vivo P. falciparum transcription samples

Comparison of transcriptional profiles of *ex-vivo* samples could reveal the genetic bases for known drug resistance and reveal new resistance mechanisms; yet the transcriptomes of parasites from these clinical samples will necessarily represent a range of different stages of progression through the IDC. A confounding factor, in this case progression through the IDC, is a variable that influences the values of another variable, in this case response to artemisinin. A confounding factor cannot be removed by changing the False Discovery Rate to zero or lowering the value used to decide the cutoff for significance because these strategies are based on the fundamental assumption that the investigator has accounted for serious confounding in the experimental design (i.e. the study compares ‘apples to apples’). Mok and colleagues used transcriptional staging to address this problem in a study of 1043 *ex-vivo* whole transcriptome samples in a study designed to detect the mechanisms for artemisinin resistance [22]. The data were aligned by comparing all single timepoint samples to a multi-timepoint reference strain (3D7) using a method similar in initial concept to the one we have employed (Methods, this paper) but without application of cyclical regression covariates.

The original Mok et al. data was run on a microarray whose probes were designed against a deprecated version of the *P. falciparum* transcriptome. Our reanalysis of this data required that we update the normalized, uncurated expression data from the Gene Expression Omnibus to a current transcriptome assembly (Methods, Additional File 1, Additional Figure 2). The transcriptional staging method used by Mok et al, applied to the recurated data for both groups confirmed the original analysis, in that Groups A and B differed in the developmental stages of parasites with Group B showing a much broader distribution (10-20hpi) than A (8-10hpi). Consequently, we predicted that the use of cyclical covariates be more impactful on Group B. To test the effectiveness of our method, we applied it to both Groups A and B; furthermore, we combined the two groups in silico (AB) to assess the effect of cyclical covariates on a sample with high variance for developmental stage.

One of the core analyses in the source paper was a correlation of gene expression values to a measure of artemisinin resistance for each sample (patient clearance half-life. We generated the differential gene expression analysis described in the Mok et al. on the recurated, transcriptionally staged but unadjusted data then compared our results for the unadjusted analysis on groups A, B and on the combined data (Group AB) with their results for groups A and B. This analysis produced results very similar to those of the original analysis (Figure 4A). We then performed the same analysis with the inclusion of cyclical regression covariates, using the inferred IDC progression time calculated by the authors and included in the metadata. We considered any genes that correlated to the artemisinin phenotype in either direction at p < 0.01 and FDR < 0.05 as significantly correlated. Comparison of these results demonstrates the effect of the cyclical regression covariates on the analysis.

**Figure 3:**
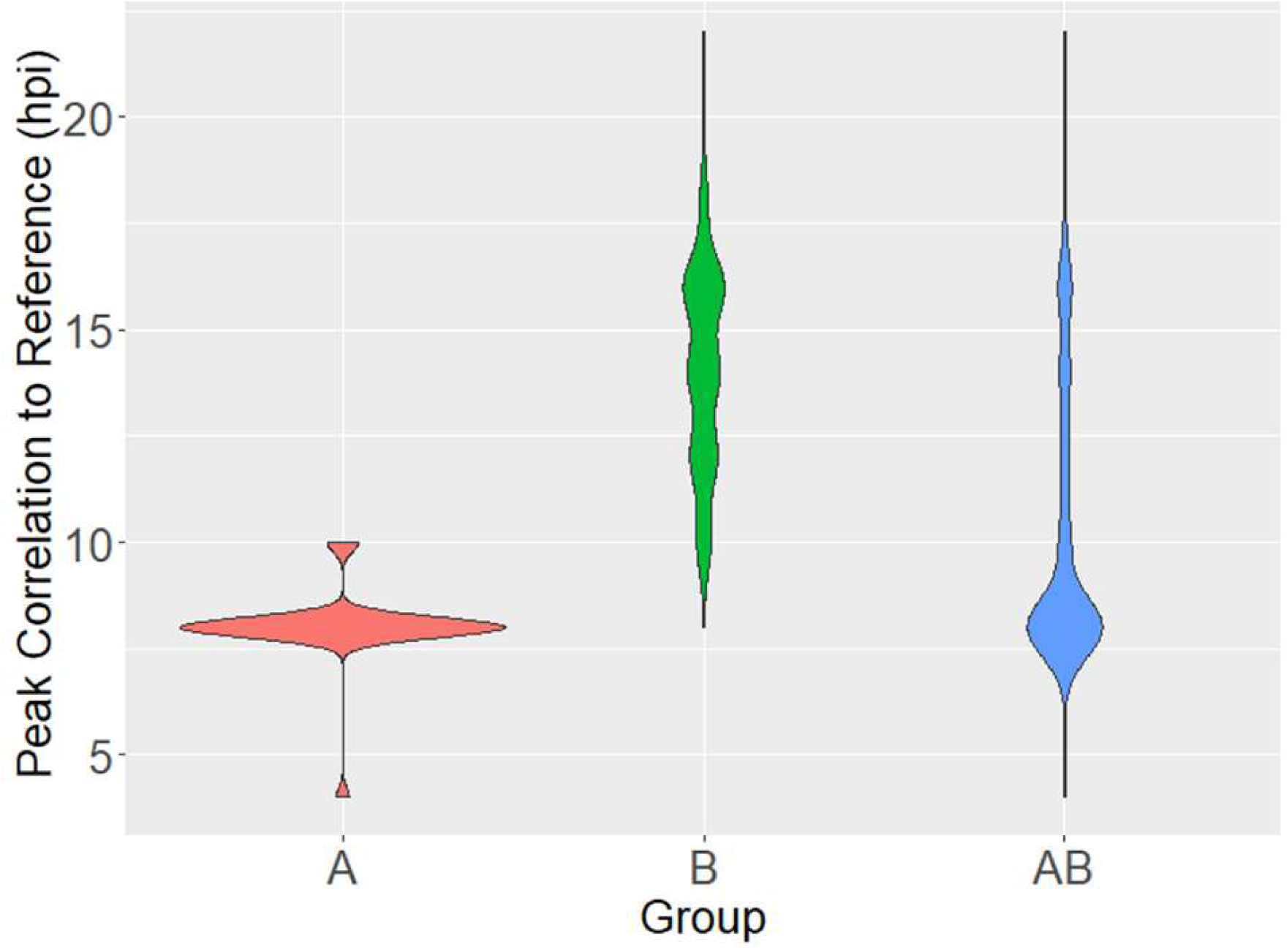
Stage Distribution of Mok et al. data. Transcriptional samples in Groups A, B, and the combined Group AB staged by transcription (Methods); displayed are the distributions of the maximum correlation of each sample to each timepoint in the reference time course in hours post invasion (hpi) [30]. The distribution in Group A is relatively narrow; the distribution in Group B and the combined Group AB are far wider.

**Figure 4:**
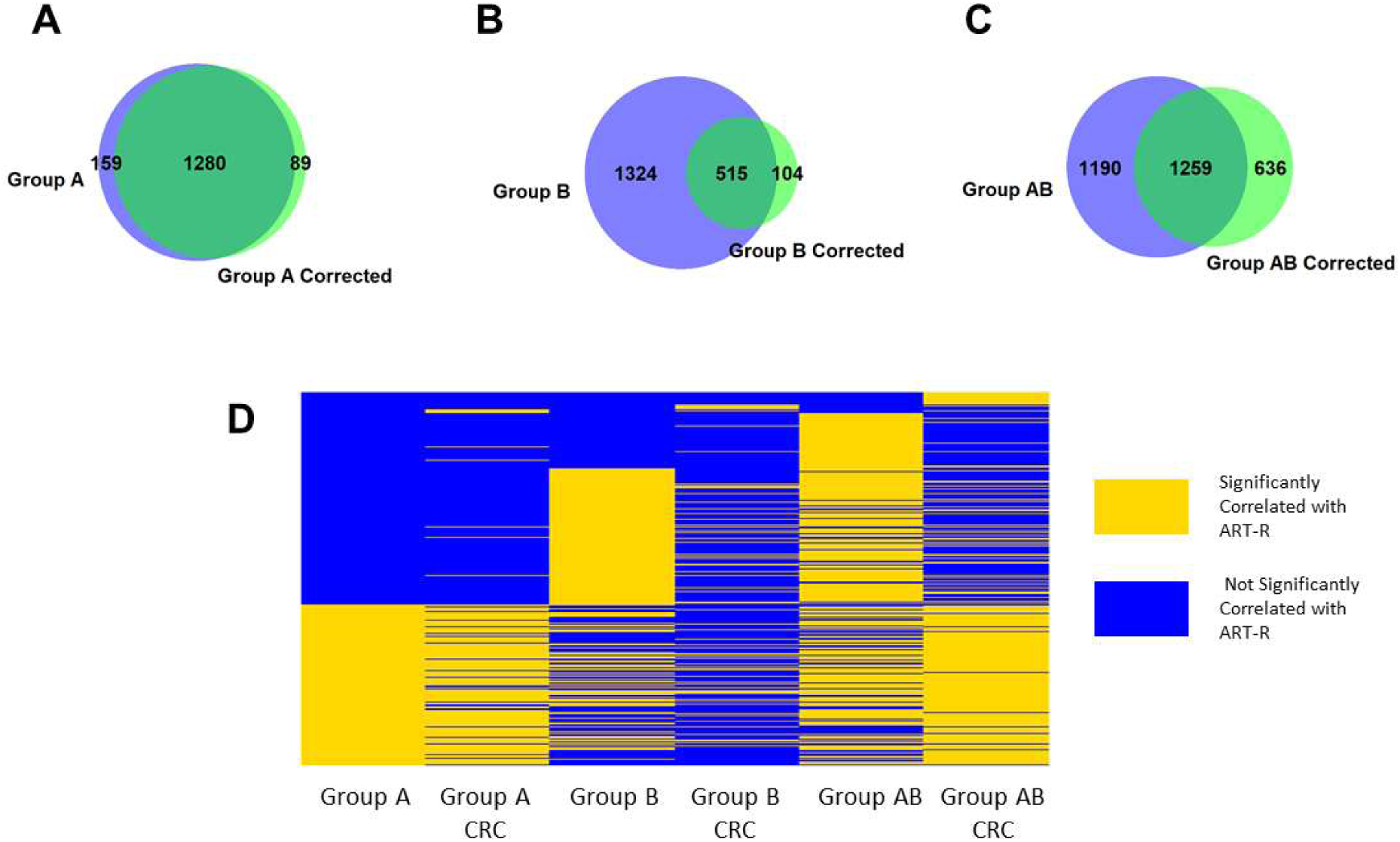
Overlap of genes considered significant before and after correction. We compared the genes considered significantly correlated with patient clearance half-life in each group before and after the application of cyclical regression covariate correction. As expected, genes considered significantly correlated with patient clearance rate (FDR < 0.05) are largely unchanged in Group A (A); in Groups B (B) and AB (C), cyclical regression covariate adjustment removes a substantial number of false positives. Figure 4D is a binary heatmap allowing for comparison between groups; each row represents a gene considered significantly correlated to artemisinin resistance (ART-R) in at least one group analysis/method combination. This allows us to observe the effect of cyclical regression covariate correction across groups. Many of the genes considered significantly correlated with patient clearance half-life in Group B uncorrected are not found in Group A; these are mostly rendered insignificant by correction. Additionally, correcting the combined Group AB recovers the majority of genes identified as significant in both groups.

Group A shows little difference in genes considered significantly correlated between the original and developmentally aligned (adjusted) analyses with an 83.8% overlap (Figure 4A). Group B shows a much larger difference, with only a 26.5% overlap (Figure 4B). The combined Group AB has a similarly small overlap, at 40.8%. Notably, the data adjusted with CRC results in fewer false positives compared to data with considerable misalignment, while few false negatives result when the data are already in good alignment; this is entirely consistent with IDC progression being observed as the single largest biological predictor of gene expression levels in *P. falciparum* [30].

If we assume genes identified as differentially expressed in group A and B after removal of the confounding factor of developmental progression are correct (i.e. no Type I error), then we can test the performance of the CRC method on the combined AB data, assuming that perfect performance will identify the same set of differentially expressed genes. Application of the Mok method to the combined data detects 875 of 1369 genes detected in the adjusted Group A (63.9%), 496 of 619 genes detected in the adjusted Group B (75.8%), and includes 749 genes not identified in either Group A or B, a potential false positive rate of 30.6%. With cyclical regression covariates, we correctly detect 1260 of 1369 genes in the adjusted Group A (92%) and 437 of 619 genes in the adjusted Group B (70.6%). Only 195 genes are present in the adjusted Group AB are not in the adjusted Groups A or B, a potential false positive rate of only 10%.

## Discussion

Alignment of the time of sampling for *P. falciparum* transcriptional experiments is critical for detection of strain-specific drug responses in malaria parasites. Alignment is technically possible with cultured *in-vitro* samples, but variance in synchronization, identification of 0hpi, IDC length and IDC progression all contribute to misalignment. Samples taken directly from the patient in a clinical setting (*ex-vivo* samples) may capture a wide range of the IDC. The clinical assessment of stage with microscopy is not sufficiently accurate to account for the developmental confounding across the entire IDC. Nevertheless, sequencing of parasite transcriptomes directly from blood, when compared to the patient’s response to ACT, permits rapid detection of emerging resistance mechanisms if the samples can be aligned to a common 0hpi. Here we demonstrated how misalignment of IDC between samples can generate both Type I and Type II errors. We further demonstrated the potentially strong impact of these errors on biological interpretation of gene expression associated with drug resistance, using published *in-vitro P. falciparum* time course data. These examples emphasize the value of a simple computational method that can disentangle expression differences that result from developmental stage in *P. falciparum* from expression differences caused by genetic differences between parasite strains.

Transcription due to developmental stage affects the majority of expressed transcripts and takes the form of an expression cascade wave that operates independently of perceived morphological stage in apicomplexan intracellular parasites [16, 30]. Setting stricter arbitrary thresholds for significance, a standard method for Type I error reduction at the price of increased Type II error, is based on the assumption that the investigator has already accounted for serious confounding in the experimental design i.e. the study compares ‘apples to apples’. When IDC progression variance is present between the samples in a single timepoint study, gene expression patterns will be driven by the IDC variance among samples. Our method for removing this confounding uses the reference time course established in multi timepoint studies in which the characteristic shape and period of the *P. falciparum* IDC transcriptional cascade was first described and characterized [17, 22, 32, 33]. We enable other investigators to apply this method with the R package **PFExpTools** posted on GitHub. In our examination of the PB58 strain from an *in-vitro* transcription experiment, we demonstrated that the use of cyclical regression covariates eliminates the false positives and the false negatives between timepoint comparisons. Application of our method to the Mok et al. data set suggested that in a set of *ex-vivo* samples in which many are in the later stages of IDC progression, the number of false positives is greatly reduced. Both false positives and false negatives are generated because the variance that drives the differences is not accounted for. Thus, the value of the arbitrary significance threshold lacks meaning because the hypothesis is predicated on the assumption that the experimental condition (drug treatment) can be fairly compared across all samples. In samples where the variance in the IDC stage is large, as in the B group, the misalignment is also large, and thus the appearance of more false positives, some of which may seem highly significant. These transcripts and the neighboring transcripts may then be subject to a rigorous but potentially misguided investigation focused on elucidating mechanisms for ACT resistance. In contrast, in Group A, in which the variance in IDC progression was lower, cyclical regression covariates had minimal impact on the result.

A recent study illustrating that artemisinin resistance mechanisms are neither as simple nor as fully elucidated as once thought serves to remind us that ACT is the last line of defense for against blood stage malaria parasites [32]. The straightforward CRC method presented here, by reducing the false positives generated by misaligned developmental stage progression, is a step toward a more precise understanding of the mechanisms of resistance artemisinin and other antimalarial drugs.

## Conclusion

Here we have demonstrated the use of cyclical regression covariates to reduce the major confounding factor in studies focused on the detection of expression difference between strains *of P. falciparum*: the unusual periodicity of transcription due to developmental progression through the IDC. Cyclical regression covariates have immediate application to studies where in vitro synchronization of all samples to the same developmental timepoint is not feasible, primarily parasite transcriptome sequencing direct from clinical blood samples, a widely used approach to frontline detection of emerging drug resistance.

## Methods

### Demonstration of the effects of progression misalignment

In order to demonstrate the effect of shifts in IDC timing on gene expression, two different data sets were used. The model dataset (Genes A and B) are purely artificial timepoints on a sinusoidal curve with a period of 48 hours. To demonstrate the effect of shifts in IDC timing in real *P. falciparum* time course data, we obtained multiple timepoint transcription data from PlasmoDB.org for two strains, 3D7 and HB3. [16, 30] The genes used as examples (PF3D7_1034400 and PF3D7_0926700) were chosen based on their representation of cyclic expression and their similarity (PF3D7_1034400) or difference (PF3D7_0926700) between strains. Unadjusted data was graphed as-is; adjusted data was plotted against the actual sampling time plus 8 hours, with any timepoint past 48 hours being plotted as the timepoint minus 48 in order to account for the 48-hour IDC.

### Determination of IDC Timing Using Transcriptional Profiles

For determination of the actual point in IDC progression for a given sample, Pearson Correlation Coefficients (PCC) between each sample’s whole transcriptome profile and the profile of each timepoint in the 3D7 reference IDC time course (hourly timepoints in the *in-vitro* 3D7 IDC) were determined.[30, 33] The hours post-infection timepoint (hpi) corresponding to the maximum PCC was considered the sample’s IDC timing, and was used in subsequent steps. This was performed using the function *stagingByTranscription()* in the R package **PFExpTools**.

### Determination of Cyclical Regression Covariates

With samples taken at different points in progression displaying differential gene expression based solely on progression, it is necessary to identify and remove any cyclic effect in a sample set. With a linear effect, we could use a single covariate related to the confounding linear effect to correct; as our confounding effect is cyclical, we require two covariates. We calculated paired cyclical regression covariates for each transcriptional sample to correct for progression’s effect on gene expression. The progression time calculated previously (T) was used in the following formula:

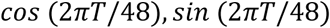

The covariates were generated using the function *CRCgeneration()* in the R package **PFExpTools**.

### Initial Application of Cyclical Regression Covariates

For testing the initial proof of concept, we looked for a dataset containing multiple replicated timepoints from a single parasite strain; this allowed us to associate any observed differences between timepoints as progression driven. The gene expression data used to demonstrate the efficacy of the cyclical regression covariate method were obtained from the Gene Expression Omnibus; the data for PB58 are listed under Accession Number GSE119514. We used *lm()* in R; for the uncorrected analysis our model was `Timepoint ∼ Gene Expression` for each gene, and for the corrected analysis the model was `Timepoint ∼ Gene Expression + xcovariate + ycovariate`. Covariates were generated as described previously. Genes were considered differentially expressed if the p-value of fit was < 0.01 and FDR < 0.05.

### Application of Cyclical Regression Covariates on Large Ex-Vivo *Dataset*

To demonstrate the efficacy of cyclical regression covariates on a larger dataset, we looked for a dataset that contained a large number of samples of varying stage. We identified the Mok et al. *ex-vivo* dataset; this set contains 1043 samples taken at various points in IDC progression. This data was obtained from the Gene Expression Omnibus, Accession Number GSE59097.

We retrieved the normalized uncurated Mok data from the Gene Expression Omnibus in order to update the curation to current transcriptome assembly. We aligned the sequence for each probe with Version 46 of the *P. falciparum* transcriptome. Probes that had an alignment with a bitscore > 130 and no secondary alignment scored over 60 were named to the gene they aligned with [34]. In the case that multiple probes aligned to a single gene, the signals were averaged. Genes with data present in greater than 80% of samples were used in the analysis. This curation was performed using the *fullCurate()* function in the R package **PFExpTools**.

To determine each gene’s correlation with the artemisinin resistance phenotype (patient clearance half-life), we used *lm()* in R; for the uncorrected analysis our model was `Phenotype ∼ Gene Expression` for each gene, and for the corrected analysis the model was `Phenotype ∼ Gene Expression + xcovariate + ycovariate`. The covariates were generated using the function *CRCgeneration()* in the R package **PFExpTools**; the IDC progression value provided in the associated metadata was used.

Data analysis and figure generation was performed in R Version 3.3.1. Venn diagrams were created using the R package **VennDiagram** [35].

## Availability of data and materials

The HB3 and 3D7 time course data is available for download at PlasmoDB.org. The PB58 data is available at the Gene Expression Omnibus under Accession Number GSE119514. The Mok et al. dataset is available at the Gene Expression Omnibus under Accession Number GSE59097. The updated curation of the Mok et al. dataset is provided as Additional File 1, with patient sample labels in the columns and feature names in the rows. A figure demonstrating a quality control comparison of our curation of the Mok et al. dataset and their reported analysis is provided as Additional File 2. The R package **PFExpTools** is freely available for download and use at https://github.com/foster-gabe/PFExpTools, including all documentation and source code under the GPL-3 license. This manuscript was prepared using Version 1.0 of **PFExpTools**.

## Declarations

### Ethics approval and consent to participate

The only human-derived data used in this study was GSE59097; the authors of this original study declared that consent was obtained appropriately for all samples.

### Consent for publication

Not Applicable

### Availability of data and materials

The HB3 and 3D7 time course data is available for download at PlasmoDB.org. The PB58 data is available at the Gene Expression Omnibus under Accession Number GSE119514. The Mok et al. dataset is available at the Gene Expression Omnibus under Accession Number GSE59097. The modern curation of the Mok et al. dataset is provided as Additional File 1. A figure demonstrating a quality control comparison of our curation of the Mok et al. dataset and their reported analysis is provided as Additional File 2. The R package **PFExpTools** is freely available for download and use at https://github.com/foster-gabe/PFExpTools, including all documentation and source code under the GPL-3 license. This manuscript was prepared using Version 1.0.0 of **PFExpTools**, available under DOI 10.5281/zenodo.3731280.

### Authors’ Contributions

Study conceived by GF. Study designed by GF, KBS, KV, JRS, MTF. Data analysis performed by GF and KBS. Manuscript prepared by GF, KV, JRS, MTF.

### Additional Files

Additional File 1 is the result of our curation of the Mok et al. data originally sourced at the Gene Expression Omnibus at Accession Number GSE59097. The dataset was curated as described in Methods using the **PFExpTools** R package. This file is a matrix of expression values, with the features in rows and samples in columns in a comma delimited file format.

Additional File 2 shows quality control comparisons between our curation and analysis of the Mok et al. data and the originally published results.

## References

1. Organization WH: World Malaria Report. In. Geneva, Switzerland: World Health Organization; 2019.

2. Duru V, Witkowski B, Menard D: Plasmodium falciparum Resistance to Artemisinin Derivatives and Piperaquine: A Major Challenge for Malaria Elimination in Cambodia. Am J Trop Med Hyg 2016, 95(6):1228–1238.

3. Ataide R, Ashley EA, Powell R, Chan JA, Malloy MJ, O’Flaherty K, Takashima E, Langer C, Tsuboi T, Dondorp AM et al: Host immunity to Plasmodium falciparum and the assessment of emerging artemisinin resistance in a multinational cohort. Proc Natl Acad Sci U S A 2017, 114(13):3515–3520.

4. Ouji M, Augereau J-M, Paloque L, Benoit-Vical F: Plasmodium falciparum resistance to artemisinin-based combination therapies: A sword of Damocles in the path toward malaria elimination. Parasite, 25.

5. Tse EG, Korsik M, Todd MH: The past, present and future of anti-malarial medicines. Malaria Journal 2019, 18(1):93.

6. Depardieu F, Podglajen I, Leclercq R, Collatz E, Courvalin P: Modes and modulations of antibiotic resistance gene expression. Clin Microbiol Rev 2007, 20(1):79–114.

7. Borst P: Genetic mechanisms of drug resistance. A review. Acta Oncol 1991, 30(1):87–105.

8. Briffotaux J, Liu S, Gicquel B: Genome-Wide Transcriptional Responses of Mycobacterium to Antibiotics. Front Microbiol 2019, 10:249.

9. Rickardson L, Fryknas M, Dhar S, Lovborg H, Gullbo J, Rydaker M, Nygren P, Gustafsson MG, Larsson R, Isaksson A: Identification of molecular mechanisms for cellular drug resistance by combining drug activity and gene expression profiles. Br J Cancer 2005, 93(4):483–492.

10. Sithara S, Crowley TM, Walder K, Aston-Mourney K: Gene expression signature: a powerful approach for drug discovery in diabetes. J Endocrinol 2017, 232(2):R131–R139.

11. Nevins JR, Potti A: Mining gene expression profiles: expression signatures as cancer phenotypes. Nat Rev Genet 2007, 8(8):601–609.

12. Vembar SS, Droll D, Scherf A: Translational regulation in blood stages of the malaria parasite Plasmodium spp.: systems-wide studies pave the way. Wiley Interdiscip Rev RNA 2016, 7(6):772–792.

13. Cho RJ, Campbell MJ, Winzeler EA, Steinmetz L, Conway A, Wodicka L, Wolfsberg TG, Gabrielian AE, Landsman D, Lockhart DJ et al: A genome-wide transcriptional analysis of the mitotic cell cycle. Mol Cell 1998, 2(1):65–73.

14. Cho RJ, Huang M, Campbell MJ, Dong H, Steinmetz L, Sapinoso L, Hampton G, Elledge SJ, Davis RW, Lockhart DJ: Transcriptional regulation and function during the human cell cycle. Nat Genet 2001, 27(1):48–54.

15. Scialdone A, Natarajan KN, Saraiva LR, Proserpio V, Teichmann SA, Stegle O, Marioni JC, Buettner F: Computational assignment of cell-cycle stage from single-cell transcriptome data. Methods 2015, 85:54–61.

16. Bozdech Z, Llinás M, Pulliam BL, Wong ED, Zhu J, DeRisi JL: The Transcriptome of the Intraerythrocytic Developmental Cycle of Plasmodium falciparum. PLoS biology 2003, 1:e5.

17. Hoo R, Zhu L, Amaladoss A, Mok S, Natalang O, Lapp SA, Hu G, Liew K, Galinski MR, Bozdech Z et al: Integrated analysis of the Plasmodium species transcriptome. EBioMedicine 2016, 7:255–266.

18. Gonzales JM, Patel JJ, Ponmee N, Jiang L, Tan A, Maher SP, Wuchty S, Rathod PK, Ferdig MT: Regulatory Hotspots in the Malaria Parasite Genome Dictate Transcriptional Variation. PLoS biology 2008, 6:e238.

19. Hu G, Cabrera A, Kono M, Mok S, Chaal BK, Haase S, Engelberg K, Cheemadan S, Spielmann T, Preiser PR et al: Transcriptional profiling of growth perturbations of the human malaria parasite Plasmodium falciparum. Nat Biotechnol 2010, 28(1):91–98.

20. Siwo GH, Tan A, Button-Simons KA, Samarakoon U, Checkley LA, Pinapati RS, Ferdig MT: Predicting functional and regulatory divergence of a drug resistance transporter gene in the human malaria parasite. BMC Genomics 2015, 16(1):115.

21. Ortiz-Ruiz A, Postigo M, Gil-Casanova S, Cuadrado D, Bautista JM, Rubio JM, Luengo-Oroz M, Linares M: Plasmodium species differentiation by non-expert on-line volunteers for remote malaria field diagnosis. Malar J 2018, 17(1):54.

22. Mok S, Ashley EA, Ferreira PE, Zhu L, Lin Z, Yeo T, Chotivanich K, Imwong M, Pukrittayakamee S, Dhorda M et al: Population transcriptomics of human malaria parasites reveals the mechanism of artemisinin resistance. Science 2015, 347:431–435.

23. Zhu L, Tripathi J, Rocamora FM, Miotto O, van der Pluijm R, Voss TS, Mok S, Kwiatkowski DP, Nosten F, Day NPJ et al: The origins of malaria artemisinin resistance defined by a genetic and transcriptomic background. Nat Commun 2018, 9(1):5158.

24. Halberg F. TYL, Johnson E.A.: Circadian System Phase — An Aspect of Temporal Morphology; Procedures and Illustrative Examples.: Springer, Berlin, Heidelberg; 1967.

25. Stolwijk AM, Straatman H, Zielhuis GA: Studying seasonality by using sine and cosine functions in regression analysis. J Epidemiol Community Health 1999, 53(4):235–238.

26. Lee G, Penataro Yori P, Paredes Olortegui M, Caulfield LE, Sack DA, Fischer-Walker C, Black RE, Kosek M: An instrument for the assessment of diarrhoeal severity based on a longitudinal community-based study. BMJ Open 2014, 4(6):e004816.

27. Rumisha SF, Smith T, Abdulla S, Masanja H, Vounatsou P: Modelling heterogeneity in malaria transmission using large sparse spatio-temporal entomological data. Glob Health Action 2014, 7:22682.

28. Wu DF, Lohrich T, Sachse A, Mundry R, Wittig RM, Calvignac-Spencer S, Deschner T, Leendertz FH: Seasonal and inter-annual variation of malaria parasite detection in wild chimpanzees. Malar J 2018, 17(1):38.

29. Goldinger A, Shakhbazov K, Henders AK, McRae AF, Montgomery GW, Powell JE: Seasonal effects on gene expression. PLoS One 2015, 10(5):e0126995.

30. Llinas M, Bozdech Z, Wong ED, Adai AT, DeRisi JL: Comparative whole genome transcriptome analysis of three Plasmodium falciparum strains. Nucleic Acids Res 2006, 34(4):1166–1173.

31. Gibbons J, Button-Simons KA, Adapa SR, Li S, Pietsch M, Zhang M, Liao X, Adams JH, Ferdig MT, Jiang RHY: Altered expression of K13 disrupts DNA replication and repair in Plasmodium falciparum. BMC Genomics 2018, 19(1):849.

32. Birnbaum J, Scharf S, Schmidt S, Jonscher E, Hoeijmakers WAM, Flemming S, Toenhake CG, Schmitt M, Sabitzki R, Bergmann B et al: A Kelch13-defined endocytosis pathway mediates artemisinin resistance in malaria parasites. Science 2020, 367(6473):51–59.

33. Mok S, Imwong M, Mackinnon MJ, Sim J, Ramadoss R, Yi P, Mayxay M, Chotivanich K, Liong K-Y, Russell B et al: Artemisinin resistance in Plasmodium falciparum is associated with an altered temporal pattern of transcription. BMC genomics 2011, 12:391.

34. Hu G, Llinás M, Li J, Preiser PR, Bozdech Z: Selection of long oligonucleotides for gene expression microarrays using weighted rank-sum strategy. BMC Bioinformatics 2007, 8:350.

35. Chen H, Boutros PC: VennDiagram: a package for the generation of highly-customizable Venn and Euler diagrams in R. BMC Bioinformatics 2011, 12:35.

