## Supplemental File 2 for "Cyclical Regression Covariates remove the major confounding effect of cyclical developmental gene expression with strain-specific drug response in the malaria parasite *Plasmodium falciparum*"

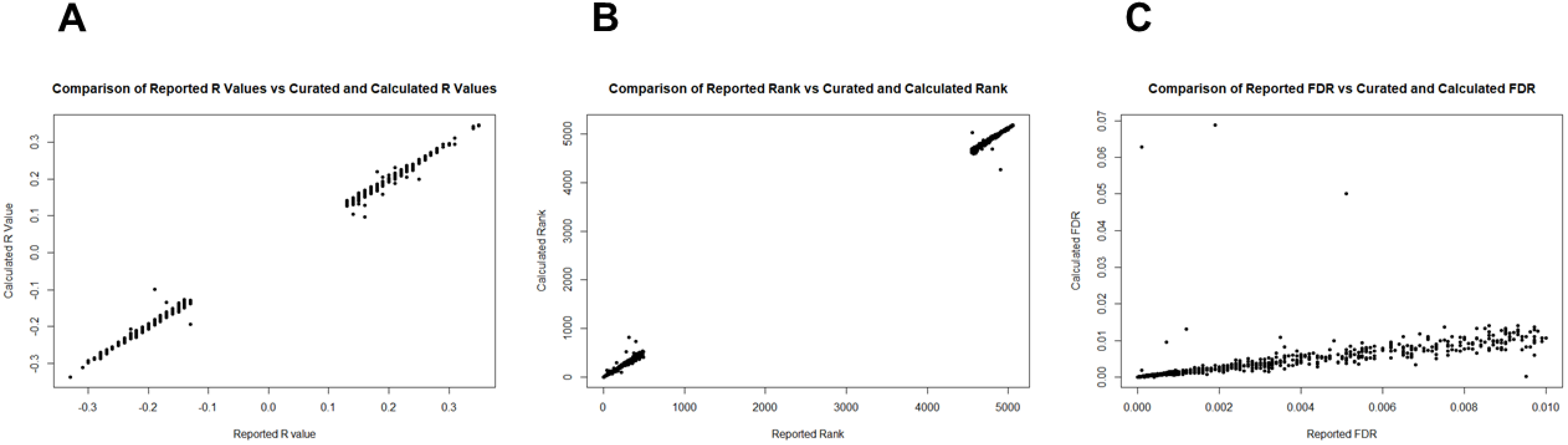


**Supplemental Figure 1: Quality Control Testing of the PFExpTools Curation Functionality.** The original Mok et al. study provided a list of genes considered significantly correlated with patient clearance half-life and associated statistics; we replicated the analysis on our curation of the probe-level data provided at the Gene Expression Omnibus. With very few exceptions, our R values (1A), correlation ranks (1B), and reported FDR values (1C) match the published results.
